# Stigmatic papilla cells recover nonself pollen acceptance after self-pollen recognition

**DOI:** 10.64898/2026.09.10.750793

**Authors:** Yuki Inoue, Sota Fujii

## Abstract

In Brassicaceae, self-incompatibility is initiated when the pollen ligand SP11/SCR activates the cognate stigma receptor kinase SRK. Although this interaction is usually considered at a single pollen-papilla interface, individual papilla cells encounter mixed pollen loads. Here, using a self-incompatibility-reconstituted Arabidopsis *thaliana* system, we show that self-pollen recognition transiently delays the hydration and germination of neighboring nonself pollen. The delay was stronger when two self-pollen grains were present but largely disappeared when nonself pollen was applied 2 h after self pollen. Repollination after removal of a self-pollen grain nevertheless revealed persistent inhibition at the original contact site. Live imaging with Lifeact-tdTomato showed that self-pollination transiently broadened F-actin orientations throughout the papilla cell, followed by recovery toward the longitudinal organization of unpollinated cells. Upon self- and non-self-dual pollination, actin-bundle orientation returned toward the papilla long axis prior to the delayed hydration of the non-self-pollen. These observations support the view that the nature of the self-recognition signaling produces a reversible, papilla-wide state that temporarily reduces compatible-pollen acceptance, while local self-rejection persists at the original contact site.

## INTRODUCTION

Self-incompatibility (SI) prevents self-fertilization by enabling the pistil to discriminate self from nonself pollen (de Nettancourt, 2001; Takayama and Isogai, 2005). In Brassicaceae, recognition specificity is determined by allele-specific binding of the pollen-coat ligand SP11/SCR to the S-locus receptor kinase SRK in stigmatic papilla cells (Schopfer et al., 1999; Takayama et al., 2000; Takasaki et al., 2000). Transgenic expression of functional S-locus components in *Arabidopsis thaliana* has provided a tractable system in which to examine the early cellular events that follow self-recognition (Nasrallah et al., 2004; Sherman-Broyles et al., 2007; Tsuchimatsu et al., 2010).

Compatible pollen rapidly obtains water and other resources from the papilla cell, hydrates, germinates, and penetrates the stigma surface. Secretory activity and a papilla-cell Ca2+-ATPase contribute to this compatible response (Iwano et al., 2014). By contrast, self-recognition blocks pollen hydration and is accompanied by a cytosolic Ca2+ signal that can spread beyond the pollen contact site (Iwano et al., 2015). F-actin in *Brassica rapa* papilla cells is also remodeled after self-pollination, and pharmacological disruption of actin compromises compatible pollen hydration and germination (Iwano et al., 2007). Together, these observations suggest that SI signaling is not confined to the receptor complex itself but can alter the physiological state of the papilla cell.

The duration and spatial range of that altered state remain poorly resolved. Under natural pollination, a papilla cell can receive self and nonself pollen in close succession. A short-term dual-pollination assay in SI-reconstituted Arabidopsis showed that self recognition could suppress a neighboring nonself grain during the first 25 min (Iwano et al., 2015), whereas classical work in *B. rapa* showed selective acceptance of nonself pollen in mixed pollen loads (Sarker et al., 1988). These findings need not be contradictory if self recognition produces a transient cell-wide response superimposed on a more persistent local response. We tested this possibility by varying the number and timing of self-pollen grains, by repollinating either the original or a distant site after removal of self pollen, and by imaging papilla-cell F-actin during self, nonself, and dual pollination.

## RESULTS AND DISCUSSION

### Self-pollen recognition transiently delays nonself-pollen hydration and germination

We placed defined combinations of pollen grains on single papilla cells expressing SRKb and followed hydration and germination by bright-field time-lapse imaging (Figure 1A); a representative mixed-load sequence is shown in Figure 1B. Pollen grains expressing SP11b were classified as self (S), whereas pollen lacking SP11b was classified as nonself (N). In the N+N condition, in which two nonself grains were placed on the same papilla, the second nonself grain hydrated after 15.4 ± 10.6 min (mean ± SD; n = 96 responding grains) and germinated after 25.2 ± 12.6 min (n = 95). When self and nonself pollen were placed in rapid succession on the same papilla (S+N), nonself hydration and germination were delayed to 64.8 ± 60.3 min (n = 66) and 72.4 ± 58.4 min (n = 63), respectively (Figures 1C–1F). The broad response-time distributions indicate that self recognition did not impose a uniform arrest; instead, it substantially extended the period before compatible-pollen acceptance.

**Figure 1.**
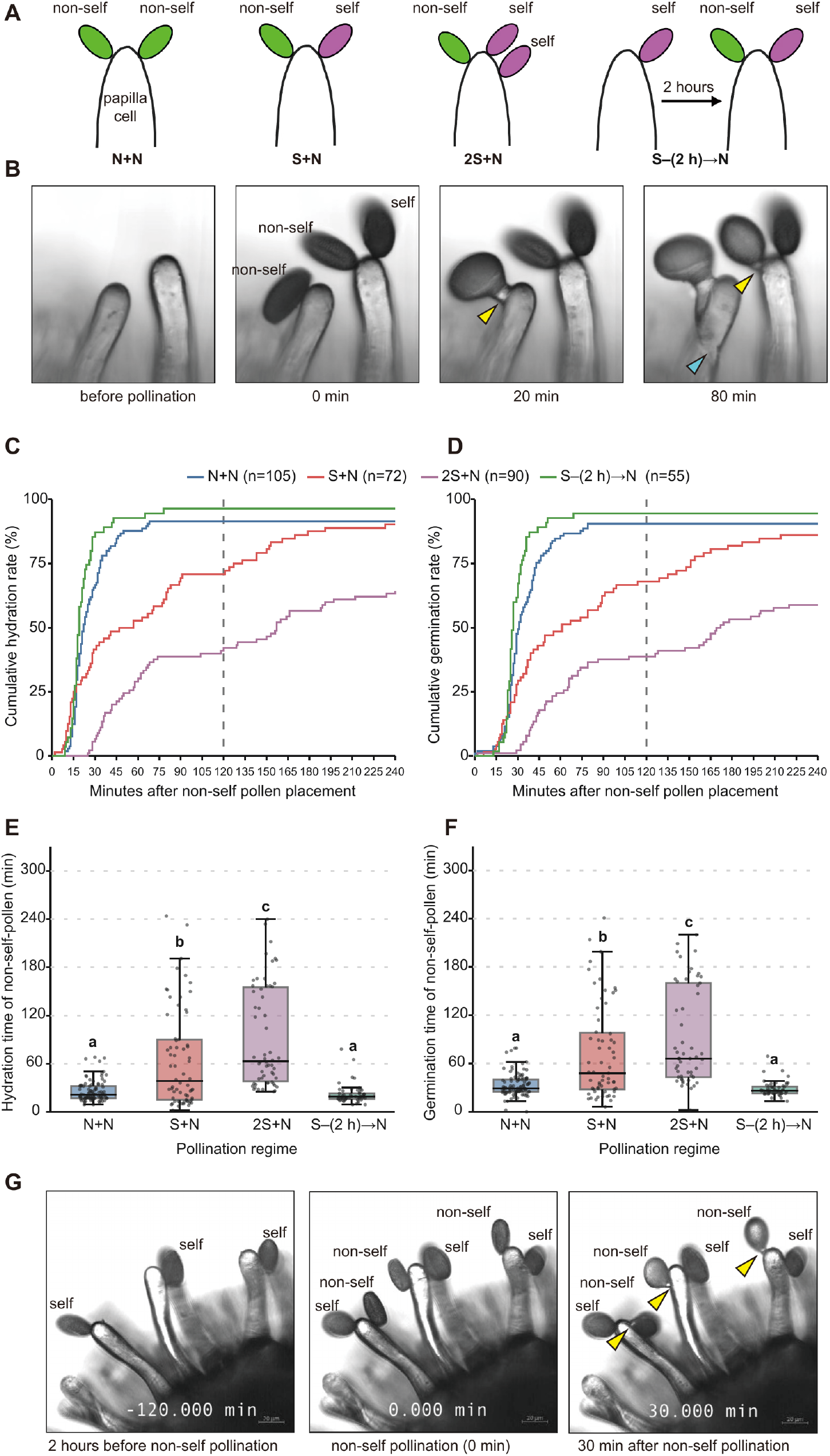
Self-pollen recognition transiently delays nonself-pollen hydration and germination. (A) Pollination regimes. N+N, two nonself-pollen grains; S+N, one self- and one nonself-pollen grain; 2S+N, two self-pollen grains and one nonself-pollen grain; S–(2 h)→N, nonself pollen placed 2 h after self pollen. (B) Representative bright-field sequence of a mixed load containing one self-pollen grain and two nonself-pollen grains. Yellow arrowheads indicate scored nonself-pollen contact sites, and the cyan arrowhead marks a pollen tube within the papilla at 80 min. (C and D) Cumulative hydration (C) and germination (D) rates from the time of nonself-pollen placement. In N+N, the second nonself grain was scored. Sample sizes indicate all scored grains: N+N, n = 105; S+N, n = 72; 2S+N, n = 90; S–(2 h)→N, n = 55. Grains without an event remained in the denominator. Dashed lines indicate 120 min. (E and F) Hydration (E) and germination (F) times for grains with observed events. Points show individual grains; boxes show medians and interquartile ranges; whiskers extend to 1.5 times the interquartile range. Different lowercase letters denote Tukey HSD groups at p < 0.05 after one-way ANOVA. (G) Representative delayed-pollination sequence showing self pollen 2 h before nonself-pollen placement (−120 min), immediately after nonself-pollen placement (0 min), and 30 min later. Yellow arrowheads mark responding nonself pollen. Scale bars in G, 20 μm.

Adding a second self-pollen grain (2S+N) further delayed nonself hydration to 84.0 ± 62.3 min (n = 58) and germination to 93.0 ± 63.0 min (n = 53). Both endpoints formed Tukey groups distinct from S+N and N+N (p < 0.05; Figures 1E and 1F), consistent with a dose-sensitive component of the papilla-wide response. The cumulative curves also show, however, that some grains did not respond within the 240-min observation period, particularly in the 2S+N condition (Figures 1C and 1D). We therefore interpret this experiment as evidence that additional self-recognition events strengthen or prolong inhibition, rather than as a precise measurement of signaling dose.

To test whether the inhibitory state decayed with time, we placed nonself pollen 2 h after self pollen [S–(2 h)→N]. Nonself grains then hydrated after 13.1 ± 12.0 min (n = 53) and germinated after 21.9 ± 12.1 min (n = 52), values assigned to the same Tukey groups as N+N (Figures 1E and 1F). The corresponding time-lapse sequence shows compatible-pollen responses while previously applied self pollen remained on the stigma (Figure 1G). Thus, the papilla-wide effect of self recognition is largely reversible over approximately 2 h. This temporal recovery reconciles the early suppression observed in dual-pollination assays with the eventual selective acceptance of compatible pollen in mixed loads.

### The self-pollen contact site retains a spatially restricted inhibitory state

Temporal recovery of whole-cell receptivity does not explain why the original self-pollen grain remains rejected. We therefore placed one self-pollen grain on a papilla, removed it after 2 h, and applied nonself pollen either to the same contact site (local) or to a different site on the same papilla (distant; Figure 2A). Only 6 of 102 nonself grains (5.9%) germinated at the local site, whereas 38 of 45 grains (84.4%) germinated at distant sites (Figure 2B). The strong spatial contrast is consistent with a long-lived, localized inhibitory state or surface modification at the original self-pollen contact site after the papilla has regained receptivity elsewhere. Together with Figure 1, it supports a two-component model comprising a reversible papilla-wide response and a persistent contact-site response.

**Figure 2.**
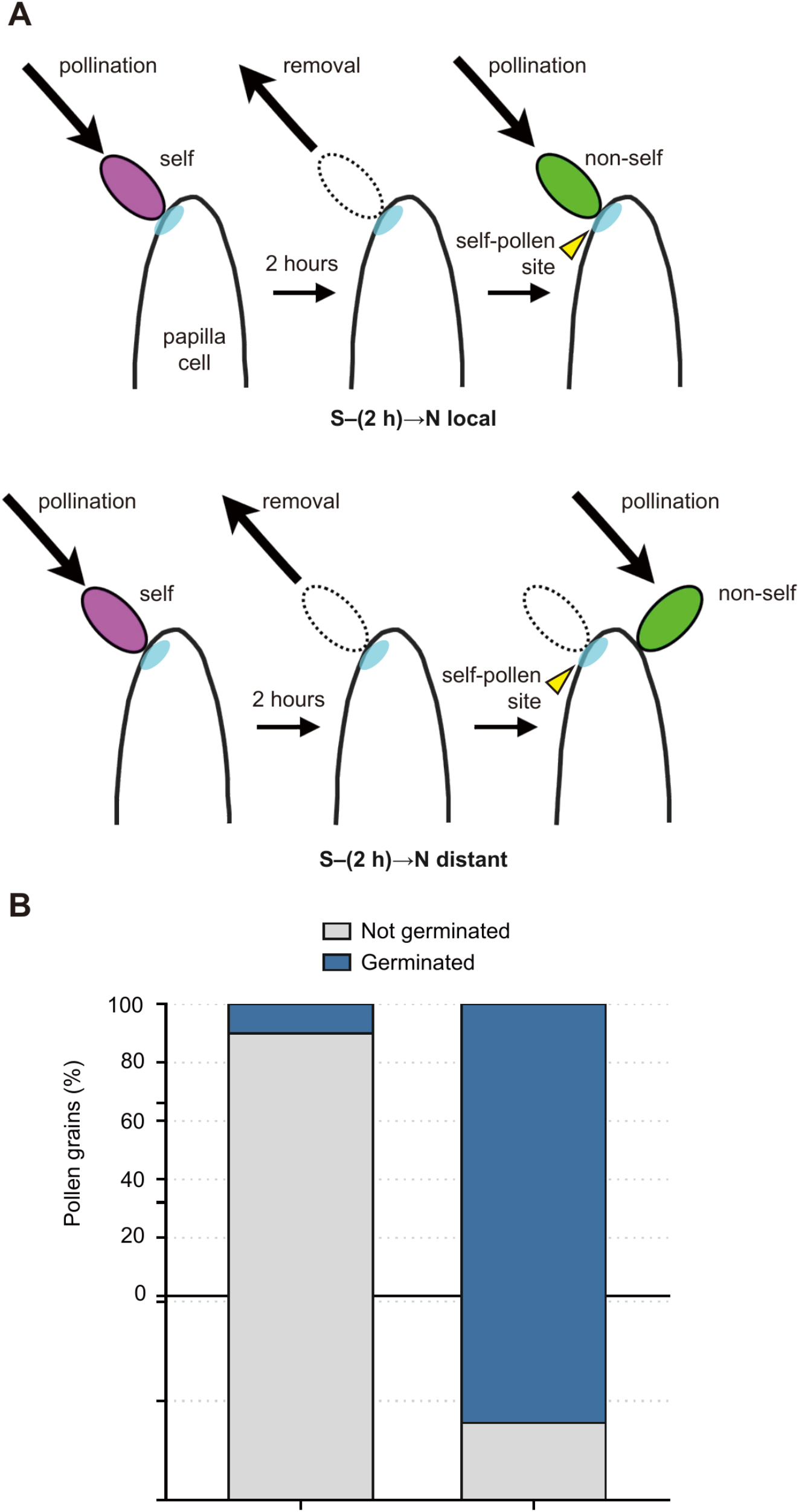
A persistent local inhibition remains at the self-pollen contact site. (A) Removal and repollination assay. A self-pollen grain was placed on a papilla and removed 2 h later. Nonself pollen was then placed either at the previous self-pollen contact site [S–(2 h)→N local] or at a spatially separate site on the same papilla [S–(2 h)→N distant]. Blue shading and yellow arrowheads indicate the original self-pollen contact site. (B) Germination outcomes shown as 100% stacked bars. In the local condition, 6 of 102 nonself-pollen grains germinated (5.9%); in the distant condition, 38 of 45 grains germinated (84.4%). Left bar, local; right bar, distant.

### Self-pollination transiently disrupts longitudinal F-actin organization

To visualize F-actin while retaining SI, we used plants coexpressing SRKb and the fluorescent F-actin reporter Lifeact-tdTomato (Era et al., 2009; Riedl et al., 2008). Aniline-blue staining confirmed abundant pollen-tube growth after nonself pollination and strong rejection after self-pollination in both the SRKb parental line and the SRKb Lifeact line (Figure 3A). The reporter therefore did not visibly abolish the SI phenotype.

**Figure 3.**
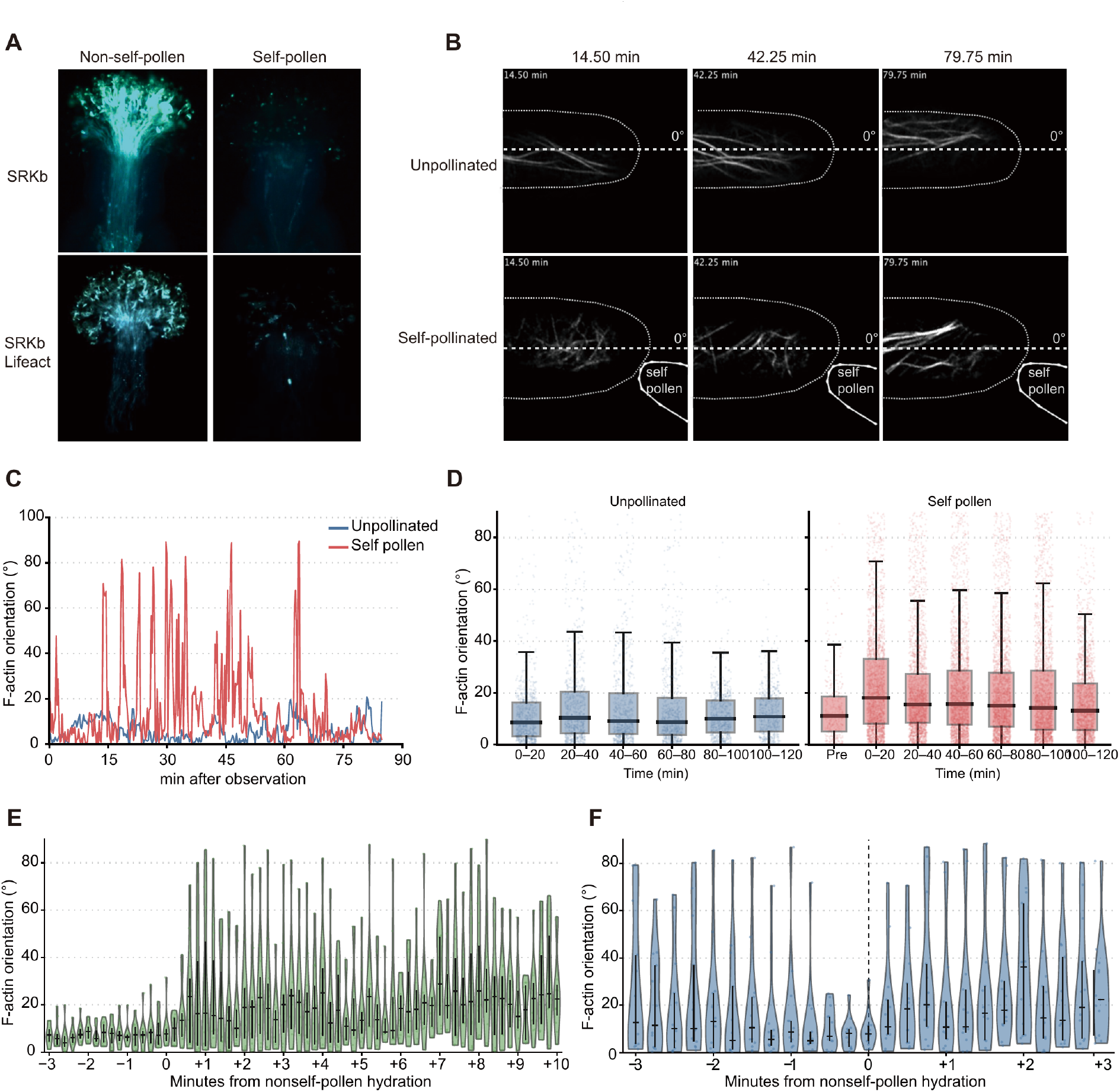
Recovery of longitudinal F-actin organization precedes delayed nonself-pollen hydration. (A) Aniline-blue staining 6 h after nonself or self pollination of stigmas expressing SRKb alone or SRKb together with Lifeact-tdTomato. (B) Representative Lifeact-tdTomato time-lapse frames from an unpollinated papilla cell and a self-pollinated papilla cell. Dotted outlines indicate cell and pollen boundaries; dashed lines mark the papilla long axis (0°). (C) Representative F-actin orientation trajectories from unpollinated and self-pollinated papilla cells. (D) F-actin orientation distributions in unpollinated cells and before (Pre) or after self-pollination, summarized in 20-min intervals. Points are image-derived orientation estimates; boxes show medians and interquartile ranges, and whiskers extend to 1.5 times the interquartile range. (E) F-actin orientation in 11 nonself-pollinated papilla-cell series aligned to nonself-pollen hydration (0 min), sampled every 0.2 min from −3 to +10 min. (F) F-actin orientation in 13 dual-pollination series aligned to hydration of the nonself-pollen grain, sampled every 0.25 min from −3 to +3 min. The dashed vertical line marks hydration onset. In E and F, violins show the distribution among tracked papilla cells, points show individual series, horizontal black lines show medians, and vertical black lines show interquartile ranges.

In unpollinated papilla cells, cortical F-actin bundles were predominantly aligned with the longitudinal axis and remained comparatively stable during imaging (Figures 3B and 3C). Self-pollination induced rapid, heterogeneous changes, including fragmentation and repeated excursions to high orientation angles in representative time courses (Figures 3B and 3C), consistent with the F-actin remodeling reported in *B. rapa* (Iwano et al., 2007). Across the image-derived angle distributions, the median orientation increased from 11.1° before self-pollination to 18.1° during the first 20 min and then declined to 13.1° at 100–120 min. Unpollinated controls remained near 9°–11° across the same interval (Figure 3D). The gradual return toward low orientation angles parallels the physiological recovery of nonself-pollen acceptance after 2 h.

### F-actin recovery precedes delayed nonself-pollen hydration during dual pollination

Compatible pollination alone produced a different temporal pattern. When nonself-pollinated series were aligned to the onset of pollen hydration, F-actin orientations were relatively low before hydration and broadened markedly during the minutes after hydration (Figure 3E). This is consistent with active cytoskeletal remodeling accompanying the transfer of water and resources to compatible pollen.

We next aligned dual-pollination series to the delayed hydration of the nonself grain. At earlier time points, orientation distributions were broad, as expected after self recognition. They became concentrated at low angles during the interval immediately preceding hydration, with the shift most evident by approximately 0.5 min before hydration, and broadened again after hydration (Figure 3F). The temporal ordering indicates that recovery of longitudinal F-actin organization is associated with renewed compatible-pollen acceptance.

The reversibility observed here differs conceptually from the gametophytic SI system of *Papaver rhoeas*, in which incompatible pollen undergoes actin-dependent programmed cell death and the inhibition becomes irreversible (Thomas and Franklin-Tong, 2004; Thomas et al., 2006). In Brassicaceae, the responding cell is the stigmatic papilla rather than the pollen tube, and recovery of papilla-cell receptivity is likely advantageous when self and nonself pollen arrive together or sequentially. Our results suggest that self recognition temporarily changes the state of the whole papilla while preserving a spatially restricted record at the self-pollen contact site.

## MATERIALS AND METHODS

### Plant material and growth conditions

*Arabidopsis thaliana* plants were grown at 22°C under a 14-h-light/10-h-dark photoperiod. SI pistils were obtained from *A. thaliana* accession C24 plants expressing the *Arabidopsis lyrata* S-b haplotype receptor kinase SRKb. Self-pollen donors were C24 transformants expressing the cognate pollen determinant SP11b; wild-type *A. thaliana* lacking SP11b served as the nonself-pollen donor. For live F-actin imaging, SRKb plants coexpressing Lifeact-tdTomato under the papilla-cell-specific AtS1 promoter (SRKb Lifeact) were used. Lifeact is a 17-amino-acid peptide derived from the *Saccharomyces cerevisiae* actin-binding protein Abp140 and labels F-actin in plant cells (Era et al., 2009; Riedl et al., 2008).

### Controlled pollination and bright-field time-lapse imaging

Flower buds were emasculated one day before anthesis. On the following day, pistils were placed on 1% (w/v) agar and secured to a coverslip with double-sided tape. Molten 1% agar was applied around tissues other than the stigmatic papillae to limit drying. Anthers were rubbed onto the coverslip, and individual pollen grains were transferred to selected papilla cells with a glass needle and micromanipulator under a stereomicroscope. Images were acquired under bright-field illumination at 1-min intervals. Time 0 was defined as placement of the scored nonself-pollen grain.

Four regimes were analyzed: two nonself grains on one papilla (N+N), with time 0 and the scored response defined by placement of the second grain; one self and one nonself grain placed in rapid succession within 5 min (S+N); two self grains and one nonself grain placed within 5 min (2S+N); and one self grain followed 2 h later by one nonself grain [S–(2 h)→N]. Hydration was scored from the visible increase in pollen-grain volume, and germination was scored at pollen-tube emergence. Cumulative response curves retained all scored grains in the denominator, including grains with no event by 240 min. Summary statistics and boxplots of event times included only grains for which the corresponding event was observed.

### Removal and repollination assay

A single self-pollen grain was placed on a papilla and removed under a stereomicroscope after 2 h. A nonself-pollen grain was then placed either at the original self-pollen contact site (local) or at a spatially separate site on the same papilla (distant). Germination was scored by bright-field observation as described above.

### Aniline-blue staining

Emasculated SRKb and SRKb Lifeact pistils were pollinated with self or nonself pollen under a stereomicroscope and incubated for 6 h at 23°C and 50% relative humidity. Pistils were fixed and cleared overnight in acetic acid:ethanol (1:3, v/v), treated with 1 N NaOH, and stained for at least 3 h at room temperature in the dark with 0.01% (w/v) aniline blue in 2% (w/v) K3PO4. Samples were mounted in 50% (v/v) glycerol and observed under UV excitation with a Carl Zeiss fluorescence microscope.

### Lifeact imaging and Airyscan processing

SRKb Lifeact pistils were prepared on coverslips as described above and imaged with an inverted LSM 880 confocal microscope equipped with Airyscan detection (Carl Zeiss). Lifeact-tdTomato was excited with a 561-nm DPSS laser, and image acquisition was controlled with ZEN 2.3 SP1 software. A Plan-Apochromat 63× oil-immersion objective was used. Time-lapse series were acquired at 1.5-to 15-s intervals. Because cortical F-actin occupied an approximately 20-μm-deep papilla volume, z-stacks were collected at 0.8-μm intervals together with transmitted-light images for pollen localization. Airyscan processing was applied in ZEN, and optical sections immediately beneath the papilla-cell surface were selected and exported as AVI files. For self-pollination experiments, an approximately 5-min pre-pollination series was acquired before self pollen was placed and imaging continued for up to 2 h. For dual pollination, self pollen was placed first and nonself pollen was added approximately 5 min later; series were subsequently aligned to the observed onset of nonself-pollen hydration.

### F-actin orientation analysis

Images were processed in Fiji (Schindelin et al., 2012) with an IJ1 macro. Frames were converted to 8-bit grayscale, cropped to the papilla-cell region of interest, band-pass filtered to reduce background and noise, and binarized with the MaxEntropy threshold. F-actin segment orientations were extracted with the LPX LinesAngle plugin (Higaki et al., 2010). The papilla long axis was measured in Fiji and defined as 0°. For each detected segment, the absolute angular difference from the papilla long axis was folded to the range 0°–90°. In Figure 3D, orientation estimates were pooled into 20-min intervals. In Figure 3E, 11 nonself-pollinated papilla-cell series were sampled every 0.2 min from −3 to +10 min relative to hydration. In Figure 3F, 13 dual-pollination series were sampled every 0.25 min from −3 to +3 min relative to nonself-pollen hydration.

### Quantification and statistical analysis

Graphs and statistical analyses were generated with R and Microsoft Excel (R Core Team, 2017). Hydration and germination times were analyzed separately by one-way analysis of variance followed by Tukey’s honestly significant difference test. Conditions assigned different lowercase letters in Figures 1E and 1F differed at p < 0.05. Boxes show medians and interquartile ranges, whiskers extend to the most extreme value within 1.5 times the interquartile range, and points show individual responding pollen grains.

## REFERENCES

de Nettancourt, D. (2001). Incompatibility and Incongruity in Wild and Cultivated Plants, Second Edition (Springer-Verlag).

Era, A., Tominaga, M., Ebine, K., Awai, C., Saito, C., Ishizaki, K., Yamato, K.T., Kohchi, T., Nakano, A., and Ueda, T. (2009). Application of Lifeact reveals F-actin dynamics in Arabidopsis thaliana and the liverwort, Marchantia polymorpha. Plant Cell Physiol. 50, 1041–1048.

Higaki, T., Kutsuna, N., Sano, T., Kondo, N., and Hasezawa, S. (2010). Quantification and cluster analysis of actin cytoskeletal structures in plant cells: role of actin bundling in stomatal movement during diurnal cycles in Arabidopsis guard cells. Plant J. 61, 156–165.

Iwano, M., Shiba, H., Matoba, K., Miwa, T., Funato, M., Entani, T., Nakayama, P., Shimosato, H., Takaoka, A., Isogai, A., et al. (2007). Actin dynamics in papilla cells of Brassica rapa during self- and cross-pollination. Plant Physiol. 144, 72–81.

Iwano, M., Igarashi, M., Tarutani, Y., Kaothien-Nakayama, P., Nakayama, H., Moriyama, H., Yakabe, R., Entani, T., Shimosato-Asano, H., Ueki, M., et al. (2014). A pollen coat-inducible autoinhibited Ca2+-ATPase expressed in stigmatic papilla cells is required for compatible pollination in the Brassicaceae. Plant Cell 26, 636–649.

Iwano, M., Ito, K., Fujii, S., Kakita, M., Asano-Shimosato, H., Igarashi, M., Kaothien-Nakayama, P., Entani, T., Kanatani, A., Takehisa, M., et al. (2015). Calcium signalling mediates self-incompatibility response in the Brassicaceae. Nat. Plants 1, 15128.

Nasrallah, M.E., Liu, P., Sherman-Broyles, S., Boggs, N.A., and Nasrallah, J.B. (2004). Natural variation in expression of self-incompatibility in Arabidopsis thaliana: implications for the evolution of selfing. Proc. Natl. Acad. Sci. USA 101, 16070–16074.

R Core Team (2017). R: A Language and Environment for Statistical Computing (R Foundation for Statistical Computing).

Riedl, J., Crevenna, A.H., Kessenbrock, K., Yu, J.H., Neukirchen, D., Bista, M., Bradke, F., Jenne, D., Holak, T.A., Werb, Z., et al. (2008). Lifeact: a versatile marker to visualize F-actin. Nat. Methods 5, 605–607.

Sarker, R.H., Elleman, C.J., and Dickinson, H.G. (1988). Control of pollen hydration in Brassica requires continued protein synthesis, and glycosylation is necessary for intraspecific incompatibility. Proc. Natl. Acad. Sci. USA 85, 4340–4344.

Schindelin, J., Arganda-Carreras, I., Frise, E., Kaynig, V., Longair, M., Pietzsch, T., Preibisch, S., Rueden, C., Saalfeld, S., Schmid, B., et al. (2012). Fiji: an open-source platform for biological-image analysis. Nat. Methods 9, 676–682.

Schopfer, C.R., Nasrallah, M.E., and Nasrallah, J.B. (1999). The male determinant of self-incompatibility in Brassica. Science 286, 1697–1700.

Sherman-Broyles, S., Boggs, N., Farkas, A., Liu, P., Vrebalov, J., Nasrallah, M.E., and Nasrallah, J.B. (2007). S locus genes and the evolution of self-fertility in Arabidopsis thaliana. Plant Cell 19, 94–106.

Takasaki, T., Hatakeyama, K., Suzuki, G., Watanabe, M., Isogai, A., and Hinata, K. (2000). The S receptor kinase determines self-incompatibility in Brassica stigma. Nature 403, 913–916.

Takayama, S., Shiba, H., Iwano, M., Shimosato, H., Che, F.S., Kai, N., Watanabe, M., Suzuki, G., Hinata, K., and Isogai, A. (2000). The pollen determinant of self-incompatibility in Brassica campestris. Proc. Natl. Acad. Sci. USA 97, 1920–1925.

Takayama, S., and Isogai, A. (2005). Self-incompatibility in plants. Annu. Rev. Plant Biol. 56, 467–489.

Thomas, S.G., and Franklin-Tong, V.E. (2004). Self-incompatibility triggers programmed cell death in Papaver pollen. Nature 429, 305–309.

Thomas, S.G., Huang, S., Li, S., Staiger, C.J., and Franklin-Tong, V.E. (2006). Actin depolymerization is sufficient to induce programmed cell death in self-incompatible pollen. J. Cell Biol. 174, 221–229.

Tsuchimatsu, T., Suwabe, K., Shimizu-Inatsugi, R., Isokawa, S., Pavlidis, P., Städler, T., Suzuki, G., Takayama, S., Watanabe, M., Shimizu, K.K., et al. (2010). Evolution of self-compatibility in Arabidopsis by a mutation in the male specificity gene. Nature 464, 1342–1346.

